# LowDoseWizard – rapid and standardised setup of low-dose cryo-TEM imaging in SerialEM

**DOI:** 10.64898/2026.05.05.722937

**Authors:** Simon A. Fromm, Simone Mattei

## Abstract

Structure elucidation of biological macromolecules by single particle cryogenic electron microscopy (SPA cryo-EM) or cryogenic electron tomography (cryo-ET) relies on low-dose imaging on cryogenic transmission electron microscopes (cryo-TEMs). Routine microscope setup remains technically demanding and can be time-consuming, particularly for inexperienced or infrequent users. We present LowDoseWizard, a guided workflow implemented in SerialEM that enables rapid and standardised setup of cryo-TEM imaging conditions. From minimal user input, the workflow configures microscope optics, camera parameters and image shift settings for all low-dose imaging states, and guides the user through key daily alignment procedures including beam shift offset calibration, objective lens astigmatism correction and coma-free alignment. The workflow is organised into modular routines that can be executed sequentially or independently, while microscope-specific acquisition parameters are defined in editable configuration files, allowing flexible adaptation to different instruments without modification of the core scripts. Across user sessions on three microscopes at EMBL Heidelberg, the complete setup required on average less than 15 minutes. To assess whether predefined imaging conditions generated by the workflow are compatible with high-resolution data collection, we acquired apoferritin data on a 200 kV Glacios and a 300 kV Titan Krios. These datasets yielded reconstructions at 1.62 Å and 1.09 Å resolution, respectively, demonstrating that rapid, guided setup can support near-atomic and atomic-resolution single particle cryo-EM. LowDoseWizard lowers the barrier to robust cryo-TEM setup, reduces the time spent on routine parameter selection and alignment, and helps users focus on sample-specific aspects of data acquisition such as target selection. The workflow should be particularly valuable in shared instrumentation environments, where accessibility, reproducibility and efficient microscope use are critical.

## 1. Introduction

Cryo-EM has become a central method for the structural analysis of biological macromolecules by single particle analysis and for the investigation of molecular organisation in situ by cryo-ET. This progress has been driven by major advances in instrumentation, including the introduction of direct electron detectors (DEDs) with improved detective quantum efficiency (DQE) and readout speed (Campbell *et al*., 2012, Li *et al*., 2013, Bai *et al*., 2013, McMullan *et al*., 2014, Ruskin *et al*., 2013, Peng *et al*., 2023), and the implementation of high-throughput acquisition strategies such as fringe-free imaging (FFI) and aberration-free image shift (AFIS) (Cheng *et al*., 2018, Eisenstein *et al*., 2023, Eisenstein *et al*., 2024). In parallel, developments in image processing have made both single particle cryo-EM (Scheres, 2012, Punjani *et al*., 2017, Kimanius *et al*., 2016) and cryo-ET (Tegunov & Cramer, 2019, Tegunov *et al*., 2021, Zivanov *et al*., 2022, Burt *et al*., 2024) increasingly robust and widely applicable. As a result, modern cryo-TEM workflows can generate very large quantities of high-quality data, placing increasing importance on acquisition procedures that are reliable, reproducible and matched to the specific experimental goal.

Despite these advances and the availability of automated sample screening and data collection software (Bouvette *et al*., 2022, Cheng *et al*., 2023), routine setup of low-dose imaging conditions on high-end cryogenic transmission electron microscopes remains technically demanding. The choice of beam conditions, magnification, fluence, defocus, camera parameters, and alignment procedures depends on both the microscope configuration and the intended application — whether screening, single particle analysis, or tomography. This challenge is particularly relevant in shared facilities, where users may have widely differing levels of experience and may spend only limited time at the microscope relative to sample preparation and downstream data processing. Commercial acquisition packages provide guided environments for routine operation, but may offer limited flexibility when workflows need to be adapted to specific specimens or when new acquisition strategies are to be implemented. By contrast, an open-source software such as SerialEM (Mastronarde, 2005) provides extensive control over microscope operation and has become a widely used platform for the development and application of advanced cryo-EM and cryo-ET workflows. Nevertheless, SerialEM’s lack of guided workflows and its open design allowing for maximal flexibility and full control of nearly all imaging parameters also requires greater user expertise during setup.

To make low-dose cryo-TEM setup faster, more reproducible and more accessible without sacrificing the flexibility of SerialEM, we developed LowDoseWizard, a guided workflow for configuring low-dose imaging sessions from minimal user input. The workflow automatically defines beam parameters, magnifications and camera settings for different imaging tasks and guides the user through key daily calibration and tuning steps. Microscope-specific settings are stored in editable configuration files, allowing adaptation to different instruments while preserving a common workflow structure. Here, we describe the design and implementation of LowDoseWizard and evaluate its use in routine cryo-TEM setup. We show that the workflow enables complete setup within approximately 15 minutes and that the resulting predefined imaging conditions support high-resolution single particle cryo-EM data collection, as demonstrated by apoferritin reconstructions at 1.62 Å on a Glacios and 1.09 Å on a Titan Krios.

## 2. LowDoseWizard design and implementation

### 2.1. Workflow overview

LowDoseWizard organises cryo-TEM low-dose setup into a sequence of modular routines that can be executed consecutively or independently, depending on the needs of the session. The workflow is typically initiated after acquisition of a grid map (also termed ‘atlas’) and proceeds from configuration of microscope optics to setting of camera parameters, followed by calibration of image shift offsets and final tuning steps such as astigmatism correction and coma-free alignment (Fig. 1). All routines are launched from a single master script, and each routine begins with a limited set of user prompts relevant to the specific imaging task (Supplementary Video S1). From these inputs, the remaining parameters are determined automatically. Manual intervention via the TEM control hand panels is required only where direct microscope interaction is necessary, and these steps are accompanied by explicit prompts within the workflow.

**Figure 1.**
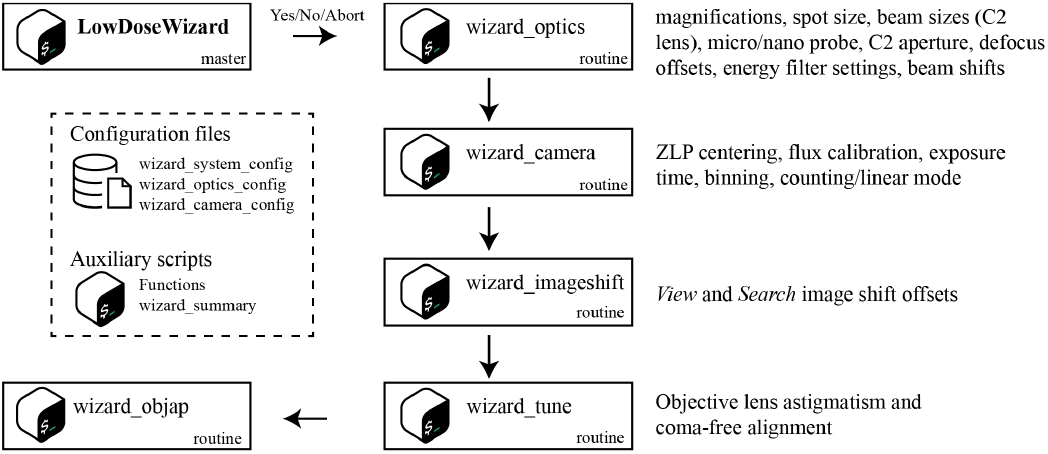
LowDoseWizard workflow.

The only prerequisite for execution is the definition in the SerialEM *Navigator* of one position over vacuum, labelled ‘empty’, and one position on a sacrificial specimen area, labelled ‘sample’. These positions provide reference locations for parameter setup and calibration. The workflow relies on the standard *Low Dose areas* defined in SerialEM (also referred to as states; SerialEM terminology is depicted in italics). *Search* is used for mapping of grid squares or lamellae, *View* for refinement of the target position following the last stage movement, *Focus* for setting the desired image defocus, and *Record* for data acquisition (Fig. 2). In tomography applications, *Trial* is kept identical to *Focus* for tracking purposes during tilt series acquisition. The workflow was developed and tested on FEI/Thermo Fisher Scientific microscopes, and microscope-specific terminology used throughout reflects this implementation context.

**Figure 2.**
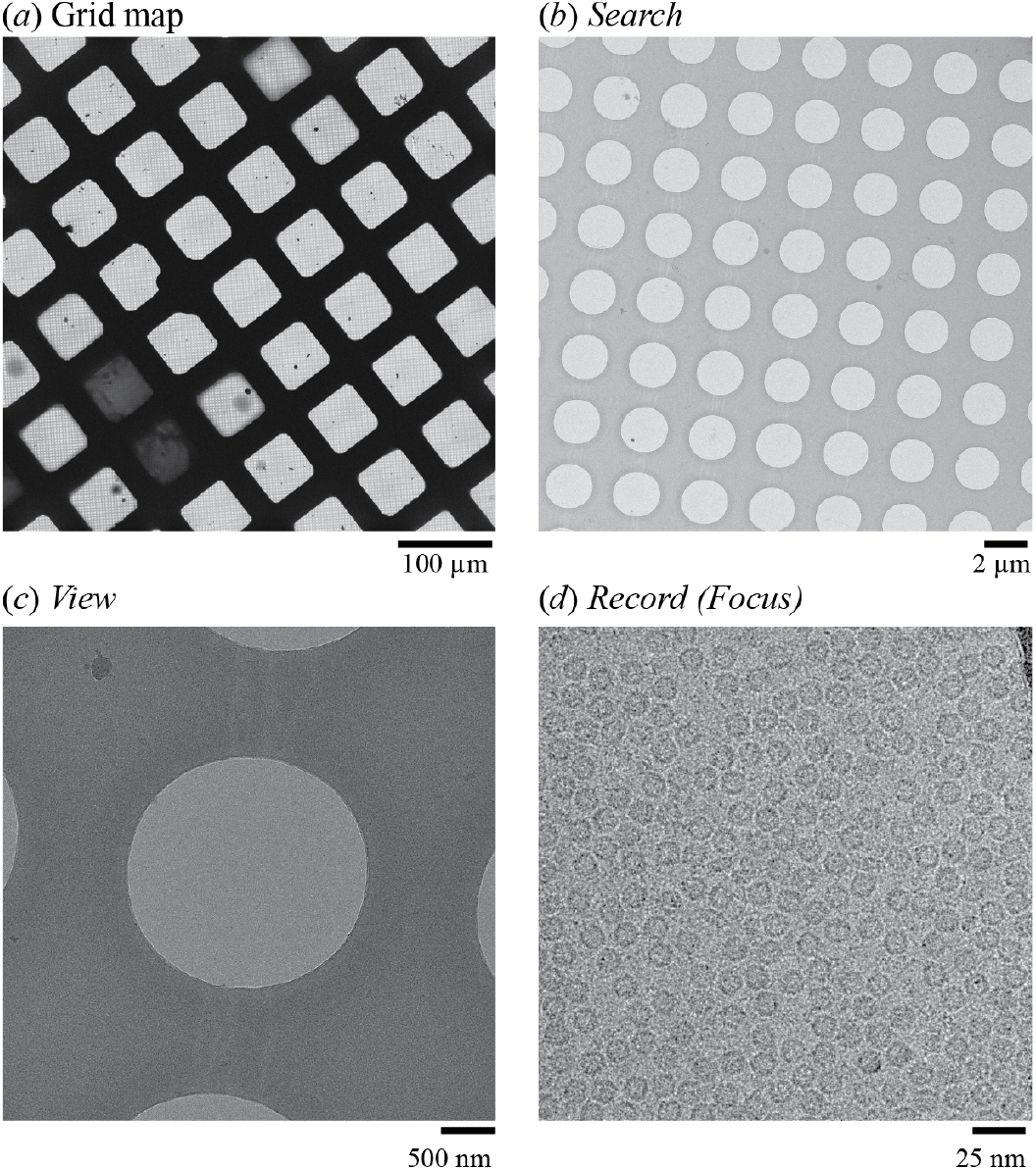
Imaging states used in the LowDoseWizard workflow. Typical field of view of an **(*a*)** grid map (100x nominal magnification, calibrated pixel size1280 Å), **(*b*)** Search (2250x, 56 Å), **(*c*)** View (11500x,11.2 Å) and **(*d*)** Record image (215000x, 0.572 Å). Images are taken from the data acquisition session on the Krios microscope.

### 2.2. Automated configuration of imaging conditions

The first stage of the workflow, named ‘optics routine’, establishes the low-dose imaging states. Because all subsequent steps depend on correctly configured *Low Dose areas*, this part of the setup has to be completed at the ‘empty’ position before calibration and tuning routines are performed. LowDoseWizard distinguishes between tomography and single particle workflows and adapts follow-up prompts accordingly. Depending on the selected application, the user specifies a small number of parameters, including the desired *Record* magnification, the required level of map detail, and, for single particle analysis, characteristics of the foil holes and support material. From these inputs, the workflow defines the appropriate settings for all *Low Dose areas*, including magnification, spot size, beam size (or C2 lens strength for two-condenser systems), C2 aperture size, minicondenser lens (microprobe or nanoprobe), defocus offsets and, where applicable, energy-filter slit width and offset. These settings are not hard-coded but are selected from microscope-specific definitions stored in a configuration file (wizard_optics_config.txt, Table 1). This allows expert users or microscope managers to establish instrument-appropriate presets once, while routine users interact only with the simplified front-end of the workflow.

**Table 1.** LowDoseWizard files.

| Script Name | File Name | Purpose |
| --- | --- | --- |
| LowDoseWizard | lowdosewizard.txt | master script: calls all routines |
| wizard_optics | 01_wizard_optics.txt | optics routine: sets magnifications and beam parameters |
| wizard_camera | 02_wizard_camera.txt | camera routine: calibrates flux and sets exposure times |
| wizard_imageshift | 03_wizard_imageshift.txt | image shift routine: aligns FOV across all low-dose states |
| wizard_tune | 04_wizard_tune.txt | tuning routine: automated astigmatism correction and coma-free alignment |
| wizard_objap | 05_wizard_objap.txt | objective aperture routine: inserts objective aperture and guides centring |
| wizard_summary | 06_wizard_summary.txt | summary routine: generates summary file |
| Functions | functions.txt | required SerialEM functions |
| n.a. | wizard_system_config.txt | configuration file for general microscope parameters |
| n.a. | wizard_optics_config.txt | configuration file for microscope optics definitions |
| n.a. | wizard_camera_config.txt | configuration file for camera parameter definitions |

The underlying parameter-selection logic is designed to preserve robust imaging conditions while adapting to different experimental goals. *Record* settings are determined directly from the user-selected magnification and the optics configuration file. Spot size is kept constant across *Low Dose areas* to minimise lens hysteresis during the session. For tomography on three-condenser systems, *View* is set to the lowest magnification that still provides a parallel beam in nanoprobe mode. For single particle applications, the *View* magnification is defined to yield a field of view (FOV) of approximately twice the user-specified foil-hole diameter, although this factor can be adjusted in the configuration file. *Search* settings likewise depend on the imaging mode: for single particle acquisition, the lowest suitable selected-area (SA) magnification is used, whereas for tomography the user can choose between minimal, medium and high detail preset levels for target selection. Where applicable, energy-filter settings are applied across all states, and for SPA sessions on gold foil *Search* can be configured for plasmon imaging (Hagen, 2022). At the end of the optics routine, the user is guided through beam shift offset calibration to ensure a centered beam in all *Low Dose areas*.

The second stage, named ‘camera routine’, configures camera parameters. Here, the user provides only the desired fluence per tilt for tomography or the desired *Record* fluence for SPA acquisition. The workflow then performs electron-flux calibration in the *Record* state at the ‘empty’ position and, if an energy filter is present, carries out zero-loss peak (ZLP) centring beforehand. From the calibrated flux, exposure times for all *Low Dose areas* are assigned automatically. As for the optics, these settings are defined through an editable camera configuration file (wizard_camera_config.txt, Table 1), which contains microscope-specific presets for quantities such as fluence, exposure time, binning and image processing. The only manual intervention required during this step is confirmation, and if necessary, correction, of beam centring in the *Record* area before flux calibration is performed. Together, these two setup stages translate a limited number of user decisions into a complete set of low-dose imaging and camera parameters tailored to the intended acquisition task.

### 2.3. Calibration and tuning routines

After the imaging states and camera conditions have been defined, LowDoseWizard performs calibration and tuning routines required for data acquisition. Image shift calibration is carried out by the ‘image shift routine’ at the ‘sample’ position for the *View* and *Search* areas. Before the calibration step, the user may interactively refine the stage position to ensure that suitable image features are present in the field of view, for example contamination, foil-hole edges or cellular structures. After eucentric height has been established, image shift offsets are calibrated automatically using the SerialEM command *FindLowDoseShiftOffset*. To guard against erroneous calibration, the resulting values are compared with predefined limits stored in the optics configuration file. If these limits are exceeded, the user is warned and can choose to repeat the calibration, keep the measured values or reset the offsets to zero.

Microscope tuning is implemented as a guided ‘tuning routine’ for objective-lens astigmatism correction and coma-free alignment. The user specifies a target defocus for tuning and may optionally request insertion of an objective aperture. Tuning can be performed either at the predefined ‘sample’ position or at the current stage position, which is useful if only the tuning part of the workflow is required. Before the automated procedure starts, the user can refine the FOV to ensure that the power spectrum contains sufficient Thon-ring signal for automated astigmatism and coma-free alignments by CTF as implemented in SerialEM. The desired defocus is then set through stage-height adjustment and, if previously inserted, the objective aperture is retracted. The automated part of the routine first corrects objective-lens astigmatism and then performs coma-free alignment. If requested, the objective aperture is subsequently inserted and manually centred, followed by a final correction of aperture-induced astigmatism. When the tuning routine is skipped, the ‘aperture routine’ for objective aperture insertion and centring can be executed independently. This grouping of calibration and tuning functions allows the workflow to accommodate both complete setup sessions and partial reconfiguration of an already prepared microscope.

### 2.4. Software architecture and configurability

LowDoseWizard is implemented entirely through the native scripting interface of SerialEM and does not require additional software beyond SerialEM itself. It uses only native script commands and is compatible with SerialEM version 4.2.2 or higher, although version 4.3.0beta or later provides the best user experience. The software architecture is deliberately modular and is divided into configuration files, scripts and reusable functions. Only the configuration files need to be adapted for a specific microscope; the scripts and functions themselves remain unchanged. This separation allows the workflow logic to be maintained independently of instrument-specific parameter definitions and facilitates transfer of the workflow between different microscopes.

Three plain-text configuration files define the microscope-specific parameters used by the workflow (Table 1). The system configuration file (wizard_system_config.txt) specifies general instrument and workflow settings, such as the presence of an energy filter, the use of a two- or three-condenser system, and the data acquisition workflows. The optics configuration file (wizard_optics_config.txt) contains microscope-specific definitions for imaging conditions, including magnifications, spot sizes, beam settings and aperture choices. The camera configuration file (wizard_camera_config.txt) defines detector-related settings such as fluence, exposure time and binning. These files are intended to be prepared once by expert staff or the microscope manager.

At the script level, LowDoseWizard consists of one master script and six routine scripts (Table 1). Only the master script is called directly by the user; it checks the required prerequisites in the *Navigator* and then invokes the selected routines in sequence (Fig. 1). Each routine may be skipped, and the full workflow may be terminated between routines if necessary. In addition, frequently reused subroutines are implemented as SerialEM *Functions*. In the current implementation, eight functions are used, including functions for low-dose area setup, camera setup, determination of eucentric height by focus and interactive stage-position refinement (Eisenstein *et al*., 2023). Because these functions are modular, they can also be called from other SerialEM scripts outside LowDoseWizard. Overall, this architecture was designed to preserve the flexibility of SerialEM while providing a standardised, guided front-end for routine cryo-TEM setup.

### 2.5. Output files and session reuse

Once the optics and camera routines have been completed, LowDoseWizard automatically generates a summary file containing the critical microscope and imaging parameters of the session and saves it in the SerialEM project folder (Supplementary Video S1). In addition to this human-readable summary, the workflow creates a session parameter file that can be used to initialise subsequent imaging sessions on the same microscope with the same parameter set. When such a parameter file is reused, the initial prompts of the optics and camera routines can be skipped. This output structure provides both a record of the configured imaging conditions and a practical mechanism for reusing validated parameter combinations across repeated sessions.

## 3. Results

### 3.1. LowDoseWizard enables rapid routine cryo-TEM setup

To assess the practical runtime of the workflow under routine operating conditions, LowDoseWizard automatically logged the duration of individual routines during user sessions on three cryo-TEMs at EMBL Heidelberg, comprising one Glacios and two Titan Krios microscopes. Timings were collected across sessions performed by 13 users, thereby capturing typical variation arising from both user interaction and microscope-specific implementation. Across all measurements, the mean cumulative runtime of the workflow was 13.5 min (Fig. 3*a*). Among the individual routines, the camera routine was the fastest and showed the smallest standard deviation, whereas the tuning routine was the slowest and displayed the largest variability (Table 2).

**Figure 3.**
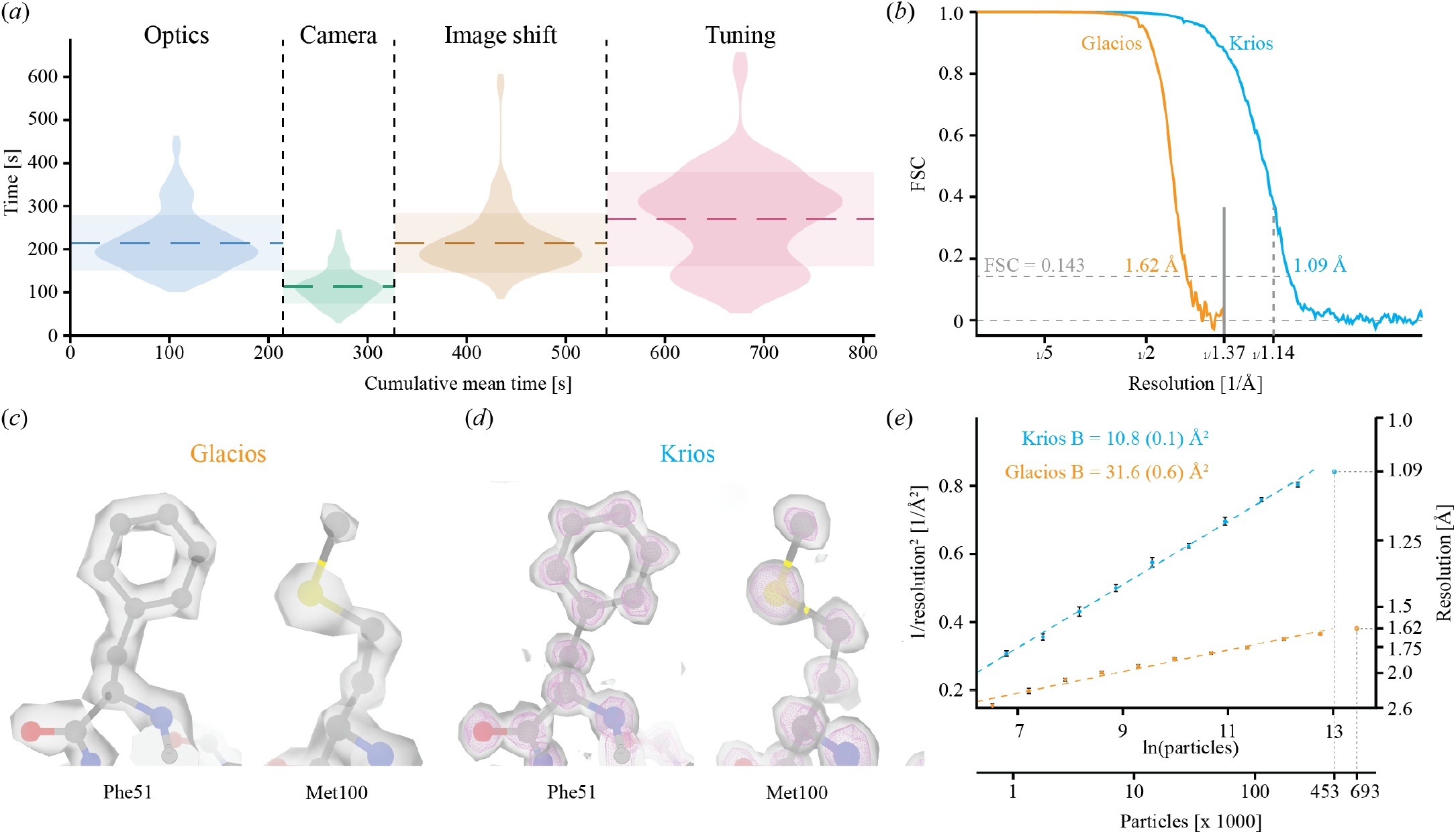
**(*a*)** Runtime of the LowDoseWizard. Dashed horizontal lines represent the mean runtime of each routine. Horizontal transparent bars represent the respective standard deviation and violine plots the raw data (Optics, n=67; Camera, n=65; Image shift, n=67; Tuning, n=45). **(*b*)** Gold-standard FSC plot of half-maps from the Krios (cyan) and Glacios (orange) data set. The grey vertical line indicates the Nyquist frequency of the Glacios data set. The dashed vertical line indicates the physical Nyquist frequency of the Krios data set. FSCs were calculated with relion_postprocess. **(*c*)** Densities from the Krios data shown at low (grey surface, level 0.2) and high (magenta mesh, level 0.52) threshold. **(*d*)** Densities from the Glacios data of same residues as in (*c*) (level 0.2). PDB 7a4m fitted into all densities. All densities were generated with relion_postprocess. **(*e*)** Rosenthal-Henderson B-factor plot (Rosenthal & Henderson, 2003) of the Krios (cyan) and Glacios (orange) data set. Error bars represent the standard deviation of the resolution estimation from the random resampling and the number in brackets the standard deviation of the fitted B-factor. The respective data point of the full particle stack (not included in the straight line fit) is highlighted by dashed lines.

**Table 2.** LowDoseWizard runtime.

| Routine | Mean [s] | SD [s] | n |
| --- | --- | --- | --- |
| Optics | 214 | 64 | 67 |
| Camera | 113 | 40 | 65 |
| Image Shift | 214 | 69 | 67 |
| Tuning | 270 | 109 | 45 |

SD: standard deviation; n: number of measurements

The limited variation in runtime of the camera routine is consistent with the small amount of user input required during this part of the workflow, which is largely determined by automated flux calibration and assignment of exposure conditions. By contrast, the larger spread observed for the tuning routine likely reflects the dependence of astigmatism correction and coma-free alignment on the availability of suitable image features and Thon-ring signal in the chosen field of view. The image shift routine similarly showed greater variability, consistent with the need for interactive refinement of stage position before calibration. Taken together, these measurements indicate that LowDoseWizard supports rapid microscope preparation in routine use, with total setup typically completed within approximately 15 min while retaining the flexibility to accommodate user- and sample-dependent adjustment during calibration and tuning.

### 3.2. Validation on Glacios and Titan Krios microscopes

To test whether the use of predefined imaging conditions generated by LowDoseWizard is compatible with high-quality cryo-EM data acquisition, we applied the workflow to benchmark SPA data collection on two microscopes at the EMBL Imaging Centre: a Glacios (Thermo Fisher Scientific) operated at 200 kV and a Titan Krios G4 (Thermo Fisher Scientific) operated at 300 kV. The Glacios was equipped with an X-FEG source, fringe-free imaging and a 20 μm C2 aperture (Konings *et al*., 2019), whereas the Krios was equipped with an E-CFEG source. Both microscopes were fitted with a Selectris X energy filter and a Falcon 4i direct electron detector.

The validation was designed to reflect realistic routine operating conditions. After loading the respective alignment file at the start of the session, no additional daily alignments were performed beyond those covered by LowDoseWizard itself. In practice, this meant that beam shift calibration, image shift offset calibration, objective-lens astigmatism correction and coma-free alignment were performed within the workflow. Of note, both TEM alignment files were last modified after major service interventions about one year prior to data collection. In addition, before starting automated data acquisition, *Coma vs Image Shift* was calibrated in SerialEM and energy-filter tuning was carried out, including zero-loss peak centring, isochromatic tuning and distortion tuning. This setup allowed assessment of whether a workflow based on predefined, microscope-specific parameter sets, supplemented by a limited number of essential session-specific calibration steps, was sufficient to support high-resolution single particle data acquisition on both a two-condenser and a three-condenser microscope.

### 3.2. Benchmark single particle data acquisition and reconstruction

Mouse heavy-chain apoferritin was used as a benchmark specimen for validation of the imaging conditions produced by LowDoseWizard. The sample was prepared at 3.2 mg ml^−1^ in 20 mM HEPES pH 7.5, 80 mM NaCl and 1.7 mM DTT. Aliquots of 3 μl were applied to glow-discharged Quantifoil R2/1 Cu 300 mesh grids carrying an additional 2 nm carbon support film. After a waiting time of 15 s, grids were blotted for 1.5 s from the sample side using a Leica EM GP2 operated at 8°C and 99% relative humidity and were plunge-frozen in liquid ethane. Data acquisition on the Titan Krios and Glacios microscopes was performed from the same grid. The principal data-collection parameters are summarised in Table 3.

**Table 3.**
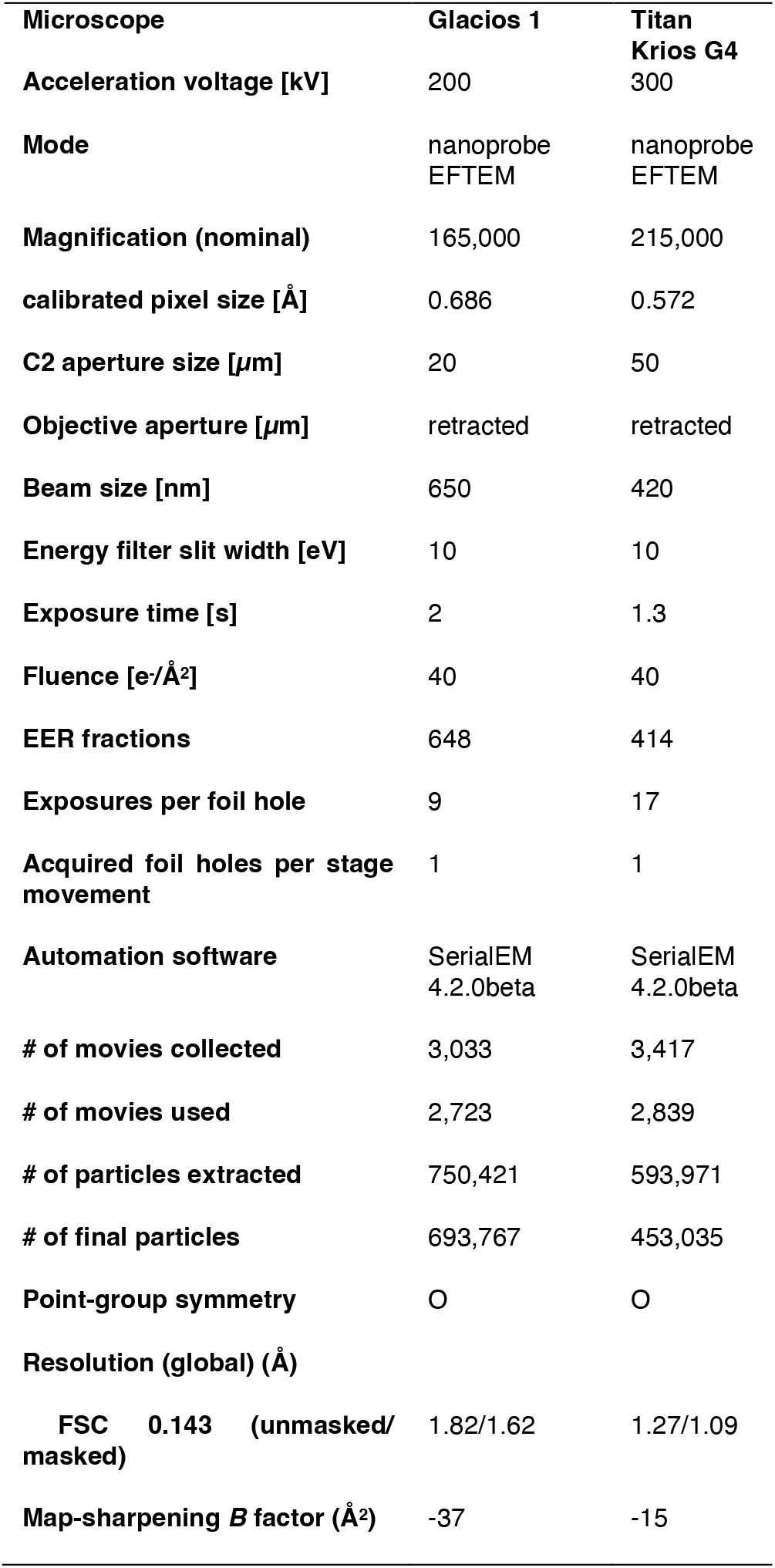
Cryo-EM data collection and reconstruction.

For the Glacios dataset, movie processing was carried out in cryoSPARC v.5.0.0 (Punjani *et al*., 2017). Movies were imported as 20 exposure groups spanning approximately 30 min intervals. After patch motion correction and patch CTF estimation, exposures with estimated astigmatism greater than 150 Å or estimated CTF resolution worse than 3 Å were discarded, leaving 2,723 micrographs for further processing. Particles were picked with crYOLO using the general pre-trained model (Wagner *et al*., 2019), yielding 750,421 extracted particles. After one round of 2D classification to remove poorly resolved particles and contaminants, 693,767 particles were retained. Refinement with octahedral symmetry, including per-particle defocus refinement, beam tilt correction, higher-order aberration refinement and Ewald-sphere correction, yielded a reconstruction at 1.72 Å resolution. One round of reference-based motion correction further improved the final masked resolution to 1.62 Å (Fig. 3*b,c*).

For the Titan Krios dataset, movies were imported into RELION 5.0 (Kimanius *et al*., 2024) and motion-corrected using RELION’s own implementation. CTF estimation was performed with CTFFIND-4.1 (Rohou & Grigorieff, 2015). After exclusion of micrographs with estimated CTF fit worse than 3 Å, astigmatism greater than 100 Å, defocus higher than −1.2 μm or CTF figure of merit below 0.2, 2,839 micrographs were retained. Particles were picked with crYOLO and classified (2D) initially in cryoSPARC v.4.7.1. After removal of poorly resolved classes, 453,035 particles were retained for refinement. Subsequent refinement with octahedral symmetry, per-particle defocus refinement, beam tilt correction, higher-order aberration refinement and Ewald-sphere correction yielded a reconstruction at 1.21 Å resolution. Bayesian polishing in RELION followed by further homogeneous refinement in cryoSPARC improved the final masked resolution to 1.15 Å, and subdivision of the dataset into six optics groups spanning approximately 60 min intervals further improved the final resolution to 1.09 Å (Fig. 3*b,d*).

The benchmark reconstructions obtained from the two datasets show that the imaging conditions generated by LowDoseWizard are compatible with high-resolution SPA cryo-EM on both microscope classes tested. The Glacios dataset yielded a final masked reconstruction at 1.62 Å resolution, whereas the Titan Krios dataset yielded a final masked reconstruction at 1.09 Å resolution (Table 3; Fig. 3*b-d*). These results indicate that rapid setup based on predefined, microscope-specific acquisition parameters does not preclude collection of data suitable for near-atomic and atomic-resolution structure determination.

To assess data quality further, Rosenthal-Henderson B factors were estimated from repeated random subsampling of the final particle stacks followed by homogeneous refinement in cryoSPARC (Nakane *et al*., 2020). The resulting B factors were 31.6 Å^2^ for the Glacios dataset and 10.8 Å^2^ for the Titan Krios dataset (Fig. 3*e*). These values are consistent with the high overall quality of both datasets and support the conclusion that, on modern cryo-TEMs, predefined imaging conditions combined with a limited number of guided calibration and tuning steps can provide acquisition conditions suitable for high-resolution SPA. Of note, high resolutions and low B factors could be achieved despite the added noise from the carbon film. Under the conditions tested here, this applied to both a 200 kV microscope equipped with fringe-free imaging and a 300 kV microscope equipped with an energy filter and E-CFEG source.

## 4. Conclusions

LowDoseWizard addresses a practical limitation of routine cryo-TEM operation by reducing setup of low-dose imaging conditions to a guided and standardised workflow implemented in SerialEM. The workflow translates a small number of user-defined choices into microscope optics and camera settings, and combines these with the calibration and tuning steps required for reliable acquisition. In routine use across multiple users and microscopes, complete setup was achieved in approximately 15 min. Validation on a 200 kV Glacios and a 300 kV Titan Krios further showed that the predefined, microscope-specific parameter sets used by the workflow are compatible with high-resolution single particle cryo-EM data acquisition, yielding apoferritin reconstructions at 1.62 and 1.09 Å resolution, respectively. Together, these results show that rapid and guided cryo-TEM setup does not compromise the quality of the resulting data under the conditions tested here.

The principal value of LowDoseWizard lies in combining the flexibility of SerialEM with a more accessible and reproducible setup procedure. Commercial acquisition environments provide strongly guided workflows for standard operation, but open platforms such as SerialEM remain important where imaging conditions need to be adapted to specific specimens or where new acquisition strategies are introduced. By structuring key setup decisions through editable configuration files and modular routines, LowDoseWizard preserves this flexibility while reducing the burden of routine parameter selection and daily alignment. This should be particularly useful in shared instrumentation environments, where users vary in experience and where microscope time is best devoted to specimen-specific tasks such as target selection. The same considerations are likely to be especially relevant for cryo-ET and correlative workflows, in which efficient setup of robust low-dose imaging conditions is only one part of a broader and often complex acquisition strategy (Nogales & Mahamid, 2024, Young & Villa, 2023).

## Acknowledgements

We thank S. Unger (EMBL Heidelberg) for valuable input to the LowDoseWizard design and critical reading of the manuscript; Z. Yang (EMBL Heidelberg) and the entire EMBL cryo-TEM user community for testing early versions of the LowDoseWizard; A. Börgel (EMBL Heidelberg) for purification of the apoferritin sample and H. Yanagisawa (Kikkawa laboratory, University of Tokyo) for the expression plasmid; D. Mastronarde for fast implementation of new commands in SerialEM; T. Hoffmann and EMBL IT Support for computational and data storage support. We acknowledge the access and services provided by the Imaging Centre at the European Molecular Biology Laboratory (EMBL IC), generously supported by the Boehringer Ingelheim Foundation.

## Data availability

All LowDoseWizard files including documentation are available on the EMBL Git (https://git.embl.org/grp-ic/lowdosewizard). Apoferritin reconstructions are available under EMDB entry EMD-57470 (Titan Krios) and EMD-57471 (Glacios). The raw EER data have been deposited under EMPIAR entry EMPIAR-13508 (Titan Krios) and EMPIAR-13509 (Glacios)

## Supporting information

Supplementary Video S1: Screen capture of the LowDoseWizard workflow on a Glacios microscope.

